# Genetic and Structural Basis of Colistin Resistance in *Klebsiella pneumoniae*: Unraveling the Molecular Mechanisms

**DOI:** 10.1101/2023.08.23.554495

**Authors:** Sahar Alousi, Jamal Saad, Balig Panossian, Rita Makhlouf, Charbel Al Khoury, Kelven Rahy, Sergio Thoumi, George F. Araj, Rony Khnayzer, Sima Tokajian

## Abstract

Antimicrobial Antimicrobial resistance (AMR), together with extensively drug resistant (XDR), mainly among Gram-negative bacteria, has been on the rise. Colistin (polymyxin E) remains one of the primary available last resorts to treat infections by XDR bacteria with the rapid emergence of global resistance. Since the exact mechanism of bacterial resistance to colistin remains unfolded, this study warranted elucidating the underlying mechanism of colistin resistance and heteroresistance among carbapenem-resistant (CR) *Klebsiella pneumoniae* isolates.

Molecular analysis was carried out on the resistant isolates using a genome-wide characterization approach, and MALDI-TOF MS for lipid A. Among the 32 CR *K. pneumoniae* isolates, three and seven isolates showed resistance and intermediate resistance, respectively, to colistin. The seven isolates with intermediate resistance exhibited the “skip-well” phenomenon, attributed to the presence of resistant subpopulations. The three isolates with full resistance to colistin showed ions using MALDI-TOF MS at m/z 1840 and 1824 representing bisphosphorylated and hexaacylated lipid A with or without hydroxylation, at position C’-2 of the fatty acyl chain, respectively. Studying the genetic environment of *mgrB* locus revealed the presence of insertion sequences that disrupted the *mgrB* locus in the three colistin resistant isolates: IS1R and IS903B. Our findings showed that colistin resistance/heteroresistance was inducible with mutations in chromosomal regulatory networks controlling lipid A moiety and IS sequences disrupting the *mgrB* gene leading to elevated MIC values and treatment failure. IS monitoring in *K. pneumoniae* could help prevent the spread of colistin resistance and decrease colistin treatment failure.

## Introduction

Antimicrobial resistance (AMR) is a global health concern (1) with predicted 10 million annual deaths worldwide by the year 2050 due to the dissemination of AMR (2). Colistin (polymyxin E) is a last-resort drug used to treat infections caused by extensively-drug resistant (XDR) Gram-negative bacteria (3). The increased use of colistin led to a rapid global spread of colistin resistance in clinically notorious Gram-negative bacteria such as *Klebsiella pneumoniae* causing both hospital and community acquired infections (4, 5). Colistin resistance could be linked to different mechanisms (6), including plasmid associated *mcr* genes (7). Intrinsic mutations, however, remain the leading cause of resistance in *K. pneumoniae* (6). The most prevalent mutations were detected in the genes encoding the PmrA/PmrB or PhoP/PhoQ two-component systems (TCSs) and in the master transmembrane regulatory protein MgrB causing changes in the LPS structure (6). These TCSs upregulate the *pmrHFIJKLM* (*pbgPE* or *arn*) operon responsible for the addition of 4-amino-4-deoxy-L-arabinose (L-Ara4N) to lipid A (8, 9). The addition of the arabinose sugar moiety increases the net negative charge of lipid A to zero and interferes with its binding affinity (10). Likewise, phosphoethanolamine (PetN) addition increases the net charge of lipid A from −1.5 to −1 (11). MgrB however, represses PhoP/PhoQ signaling (12). Insertional inactivation, missense mutations, and alterations in *mgrB* could all be also linked to high levels of colistin resistance (13). Moreover, *crrAB*, another TCS, are also associated with colistin tolerance. Mutations in *crrB*, the sensor kinase, increased the transcription of *crrC* which regulates the expression of the *pmrC* and *pmrHFIJKLM* operon through the activation of *pmrAB* (14). The emergence of heteroresistance to colistin is a growing concern as it can lead to treatment failure and the spread of resistant strains (15). Heteroresistance to colistin has significantly increased in Gram-negative bacteria (16).

Heteroresistance is defined as subsets of an isogenic bacterial population that display a range of susceptibilities to a specific antibiotic (17). The term heteroresistance also denotes genetic changes observed in subpopulations of bacteria acquiring resistance to antimicrobials (18). While heteroresistance often results in therapeutic failure (17), its detection remains difficult using standard laboratory methods (18). In *K. pneumoniae*, colistin heteroresistance has been widely documented and linked to various SNPs or gene duplications in TCSs associated with colistin resistance (19, 20). Polymyxins are cationic and diffuse poorly into agar media and susceptibility can be difficult to detect. Different dilution methods have emerged to determine the minimum inhibitory concentration (MIC) of colistin. These methods include broth microdilution (BMD) and broth macrodilution methods (6). BMD remains the gold standard and is recommended by Clinical and Laboratory Standards Institute (CLSI) and the European Committee on Antimicrobial Susceptibility Testing (EUCAST) (21, 22).

Currently, carbapenems and colistin are considered as the last resort antibiotics to combat *K. pneumoniae* related infections (23). Colistin-resistant *K. pneumoniae* (CRKP) were detected in Asia, the Americas, Europe, and in Lebanon (24-30). In this work, we aimed at elucidating the underlying mechanisms of colistin resistance in three colistin resistant and seven heteroresistant *K. pneumoniae* using a genome-wide characterization approach and MALDI-TOF MS analysis of lipid A.

## Materials and Methods

### Antibiotic susceptibility testing

A total of 32 carbapenem resistant *K. pneumoniae* clinical isolates were recovered at the Clinical Microbiology Laboratory at the American University of Beirut Medical Center (AUBMC), designated as KP1-KP32 identified by MALDI-TOF, their routine antimicrobial susceptibility testing and molecular characterization were done as reported from our laboratory by Arabaghian et al. (31). In this current study, colistin minimum inhibitory concentrations (MICs) for KP1-KP32 was determined using microdilution (BMD), population analysis profiling (PAP), E-test, and disk diffusion method with the cutoff values used being the same as recommended by Galani et al (32). Heteroresistant subpopulations were isolated through BMD and PAP, then further confirmed through E-test.

### Broth microdilution (BMD)

BMD was performed as recommended by CLSI 2018 (34) and according to the ISO standard method (20776-1). Briefly, a stock solution of 2 mg/mL colistin sulfate (Sigma-Aldrich Merck KGaA, Germany; 100 mg, 15,000 U/mg) was prepared and aliquots were stored at −80 °C. CAMH broth was purchased as ready to use Mueller-Hinton II medium (Sigma-Aldrich; Merck KGaA, Germany). Sterile non-treated polystyrene round-bottom 96-well microplates were used. Different colistin concentrations were prepared from the original stock (range, 64 μg/mL to 0.128 μg/mL in 2-fold dilution) in CAMH broth. 50 μL of 2-fold diluted colistin was dispensed in triplicates. The isolates were grown on LB agar for 24 h at 37 °C and one colony was inoculated in a 5 mL CAMH broth and adjusted to 0.5 McFarland. The culture was then diluted to obtain a final count of 5×10^5^ CFU/mL. 50 μL of the adjusted bacterial suspension was added to the wells having different colistin dilutions. A volume of 100 μL from each overnight cultured *K. pneumoniae* was loaded on the last column of each plate, preceded by a blank well containing only CAMH broth. The microplates were incubated under static conditions at 37 °C for 20-24 h. Using a microplate reader, the OD_595_ was measured after 1 and 24 h. The MIC was determined as the lowest concentration that fully inhibits bacterial growth. According to the EUCAST guidelines, isolates with MICs of ≤ 2 μg/mL are categorized as susceptible, and those with MICs of > 2 μg/mL as resistant (EUCAST, 2017). *K. pneumoniae* ATCC700603 and *K. pneumoniae* KP30 colistin sensitive isolates, were used as negative controls. Following the BMD incubation, 1 µL from each well representing all different colistin dilutions was inoculated on a colistin free agar medium. The plates were incubated at 37 °C for 24h. The obtained colonies were recovered using colistin free CAMH broth after 24 h of incubation at 37 °C and stored at −20 °C for further testing.

### E-test

Preliminary colistin susceptibility screening for KP5, KP6, KP15, KP16, KP17, KP18, KP24, KP27, KP28, KP29, KP30 was performed according to the CLSI 2017 guidelines (33). Cation Adjusted Mueller-Hinton (CAMH) agar plates (Bio-Rad, Marnes-la-Coquette, France) were used. A starting cell suspension corresponding to 0.5 McFarland was inoculated on CAMH. E-test strips (AB bioMerieux, La Balme-les-Grottes, France) were then applied, and plates were incubated for 24 h at 37 °C. BMD assay was also followed by susceptibility determination using the E-test.

### Population analysis profiling

Heteroresistance was assessed using population analysis profiling (PAP). Plating of 50 µL of bacterial cultures with an OD_595_ of 0.08-0.1 (corresponding to 0.5 Mc-Farland) was performed on cation-adjusted MH (CAMH) agar (Merck KGaA, Germany) with colistin sulphate. CAMH agar plates were prepared to have two folds increasing concentrations of colistin sulphate (1, 2, 4, 8, 16, 32, 64, and 128 mg/L). After 24 h of incubation at 37 °C the number of colonies were counted and heteroresistance assessed (35). The PAP assay was performed on colonies after their first exposure to colistin through the BMD assay. Following PAP assay, E-test method was used to determine colistin MIC. Colonies growing on PAP plates with the highest colistin concentration were sub-cultured in CAMH broth and used later to perform the E-test. The heteroresistant populations, showing the highest MICs, emerging after Post-PAP E-test were collected and further characterized.

### DNA preparation and genome sequencing

DNA was extracted and quantified by a Qubit assay using the high sensitivity kit (Life technologies, Carlsbad, CA, USA) and 0.2 µg/µL of DNA was sequenced by Illumina MiSeq runs (Illumina Inc., San Diego, USA) using Nextera XT DNA library preparation kit. The DNA was fragmented and amplified by limited PCR (12 cycles), introducing dual-index barcodes and sequencing adapters. After purification on AMPure XP beads (Beckman Coulter Inc, Fullerton, CA, USA), the libraries were normalized and pooled for sequencing on the MiSeq. Paired-end sequencing and automated cluster generation with dual indexed 2×250-bp reads were performed.

### Analysis of the *mgrB* locus and Two-component regulatory system (TCS) in *K. pneumoniae*

The PCR amplification of the *mgrB* was done using primers that cover both the *mgrB* coding and flanking regions. The primers were designated as *mgrB*_ext_F and *mgrB*_ext_R as described by (31). Sanger sequencing was performed on ABI 3500 DNA analyzer using BigDye Terminator v3.1 - Cycle Sequencing Kit (Applied Biosystems Foster City, CA, USA). The resulting gene sequence was compared with the *mgrB* sequence from *K. pneumoniae* MGH 78578 deposited on NCBI (accession number NC_009648). TCS protein sequences for (pmrA, pmrB, pmrD, phoP, and phoQ) were analyzed using PROVEAN tool (32) using *K. pneumoniae* MGH 78578 amino acid sequences as a reference. Amino acid substitutions, and indels were categorized as neutral or deleterious based on their predicted effect on the resulting protein function.

### Genome analysis

Output sequencing reads of 32 samples (KP1-32) (33) were quality checked using FastQC (34) and filtered, demultiplexed, and trimmed using Trimmomatic (35) The de novo assemblies were generated using SPAdes (36) with –careful and read error correction turned on. The resulting assemblies were checked for contiguity, completeness, contamination, and depth of coverage using QUAST (37), kraken2 (38), and Bandage (39) for manual curation of the assemblies. To detect the cationic antimicrobial peptides (CAMPs) resistance genes, we employed multiple methods. As an initial screening step, we utilized the Meta-marc tool (database model 3) (40) on assembled genomes to identify the contigs carrying cationic antimicrobial peptides (CAMPs) resistance genes. To ensure the accuracy of the results obtained from Meta-marc and to obtain the actual sequences for each identified gene in the samples, a combination of four analysis methods was used: Meta-marc, Prokka (41), Blastp against the NCBI database, and blast against the CARD (42) and MEGARes databases (43). Contigs containing CAMPs resistance genes were annotated using Prokka. The resulting nucleotide sequence file (ffn) was then used to conduct a targeted search for the resistance gene using Meta-marc. The sequences corresponding to each resistance gene were retrieved and verified by performing blastp against the NCBI and CARD databases. The obtained and validated gene sequences were aligned using MAFFT (44) and constructed a phylogenetic tree using PhyML 3.3_1 (45).

### Lipid A extraction

Lipid A extraction from all the isolates was performed on colonies obtained from the heterogeneous population without any further exposure to colistin or enrichment of the heteroresistant subpopulations. A commercial preparation (1 mg) of monophosphoryl lipid A detoxified (MPLA) from *Salmonella enterica* serovar Minnesota (R595) with a molecular weight of 1735.416 was purchased from Avanti Polar Lipids (Alabaster, AL, USA) and used as lipid A standard. Final concentrations of 1000 ppm and 100 ppm were prepared from the initial 1 mg powder stock by dissolving it in chloroform-methanol (1:1, v/v). Spotting on MALDI-TOF MS plate was done through both layering and mixing approaches. First, 0.4 μL of lipid A standard was spotted on MALDI plate in three layers and was allowed to air dry, then layered with 0.4 μL of 20 mg/mL norharmane matrix in chloroform-methanol (2:1, v/v) (Acros, New Jersey, USA).

Using the Applied Biosystems SCIEX 4800 MALDI TOF/TOF MS/MS (Applied Biosystems, Inc., MA) the signal was detected while operating in the negative reflector mode. Lipid A was extracted from *K. pneumoniae* following, with modifications, Caroff et al (46). Briefly, lyophilized phenol-inactivated *K. pneumoniae* with a mass of 30 mg was used in the acetic acid hydrolysis reaction. 400 μL of 1% acetic acid were added over the sample, followed by boiling at 100 °C in a shaking thermomixer for 1 h. To separate cell debris and retain the supernatant, the sample was cooled using an ice bath for 10-15 min and centrifuged at 2,000 xg for 15 min. The supernatant was transferred to a new tube and diluted with HPLC grade water in a ratio of (1:1, v/v), and subjected to lyophilization. The resulting pellet was purified following an optimized procedure. Briefly, the lyophilized pellet was washed twice with 400 μL of HPLC grade methanol to remove residual contaminants. Occasional vortexing and sonication for 5 min was necessary to disrupt the pellet. Pelleting was done at 8,000 xg for 5 min and after each wash reconstituted in 75 μL chloroform-methanol (2:1, v/v), vortexed, and sonicated for 5 min at 50 °C. Dowex 50W-X8 H^+^ ion exchange resin (Acros, New Jersey, USA) was added in small amounts to the mixture and vortexed. 0.4 µL of lipid A suspended in chloroform-methanol (2:1, v/v) was applied in three layers on MALDI-TOF plate. Various volumes of matrix aliquots were used to cover the sample (Figure 1). The ion acceleration voltage was set to 20 kV. Extraction delay time was adjusted within 100 to 300 ns to obtain the optimal spectra.

**Figure 1:**
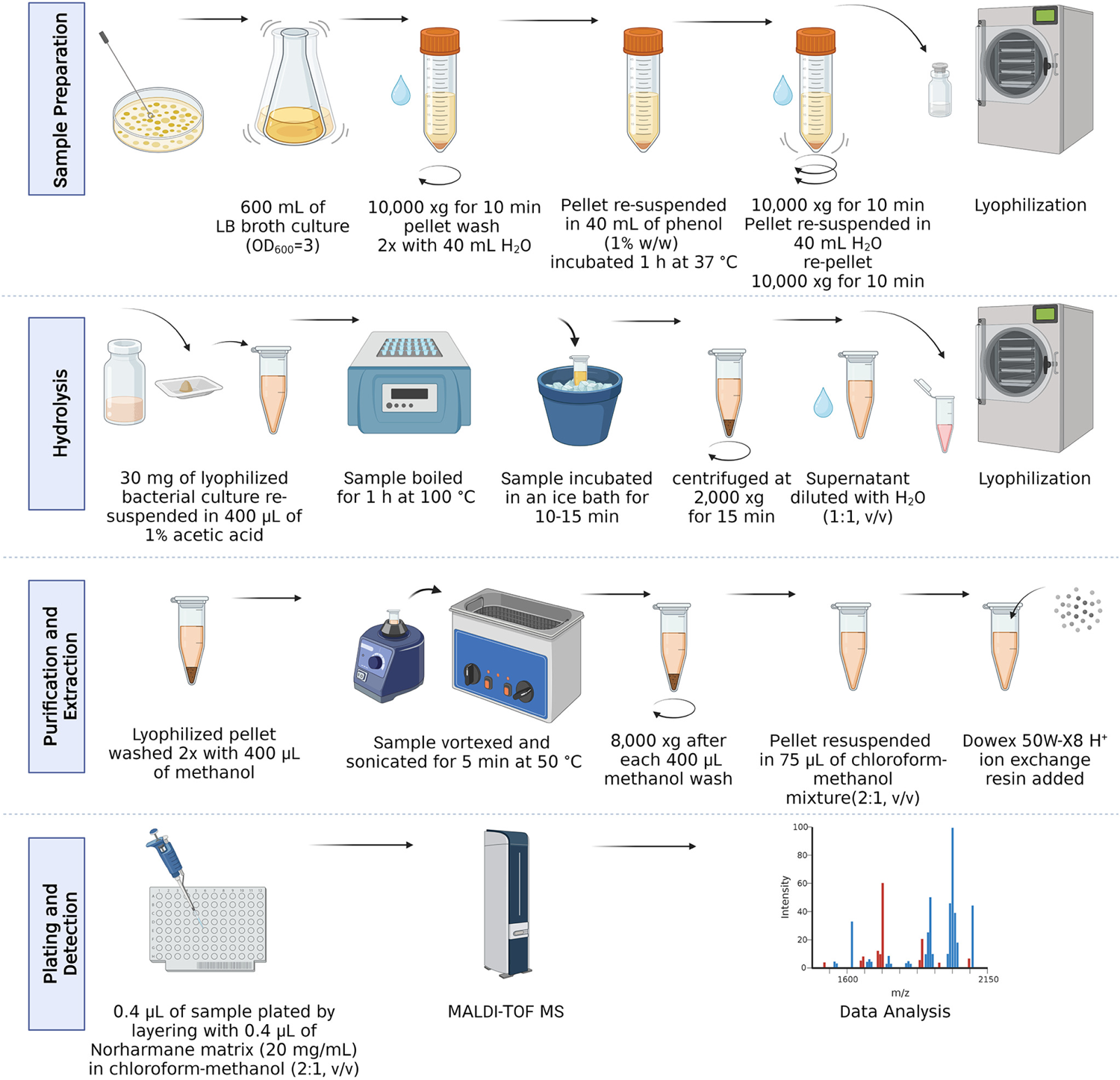
Illustration representing Lipid A extraction procedure. *K. pneumoniae* sample preparation, hydrolysis reaction, purification and extraction, and plating procedures are summarized to reflect the workflow to obtain Lipid A mass spectra.

## Results

The complete resistance profile and metadata for the isolates showing resistance/intermediate resistance (KP5, KP6, KP15, KP16, KP17, KP18, KP24, KP27, KP28, KP29, KP30) in this study are shown in (Figure 2).

**Figure 2:**
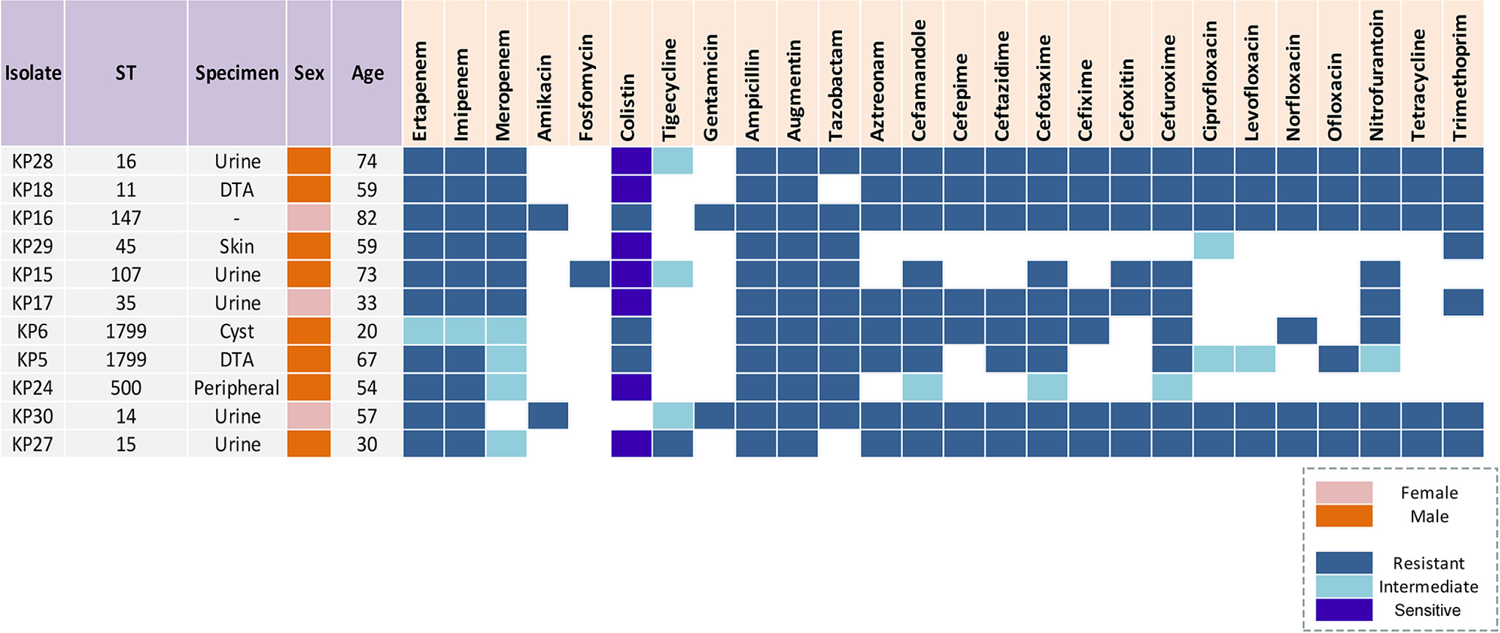
Resistance profile of *K. pneumoniae* isolates. Preliminary colistin resistance profile is shown based on routine disk diffusion results. Metadata including (ST: sequence type, Specimen: isolation site, sex, and age).

### Colistin antimicrobial susceptibility testing

Most of the tested isolates (10/11) were resistant to at least one of the three used carbapenems in this study (imipenem, meropenem, and ertapenem), except for KP6 which showed intermediate resistance to all (33). E-test results did not show any phenotypic heterogeneity towards colistin, as no discrete colonies were observed in the inhibition zone (Figure 3-A and 4-A). E-test antimicrobial suitability testing (AST) results were, however, inconsistent when compared to BMD AST. Most of the isolates, except KP5 with MIC 6 µg/mL, KP6 with MIC 4 µg/mL, and KP16 with MIC 24 µg/mL, were found to be susceptible to colistin through E-test screening (Figure 3-A, and Figure 4-A). BMD results showed higher MIC values than that obtained from the E-test. KP5 and KP6 showed MIC values of > 64 μg/mL and KP16 = 64 μg/mL (Figure 5-A). Colistin phenotypic resistance profile was also determined through disk diffusion assay, which showed that KP5, KP6, and KP16 were resistant. It is worth noting that KP15, KP17, KP18, KP24, KP27, KP28, and KP29 exhibited the “skip-well” phenomenon which could be attributed to the presence of resistant subpopulations (Figure 5-B). The test was repeated several times to rule out experimental errors, and each isolate was tested in an individual 96-well plate. *K. pneumoniae* ATCC 700603, a colistin susceptible and a negative control reference strain, showed MIC of 0.5 μg/mL using BMD assay.

**Figure 3:**
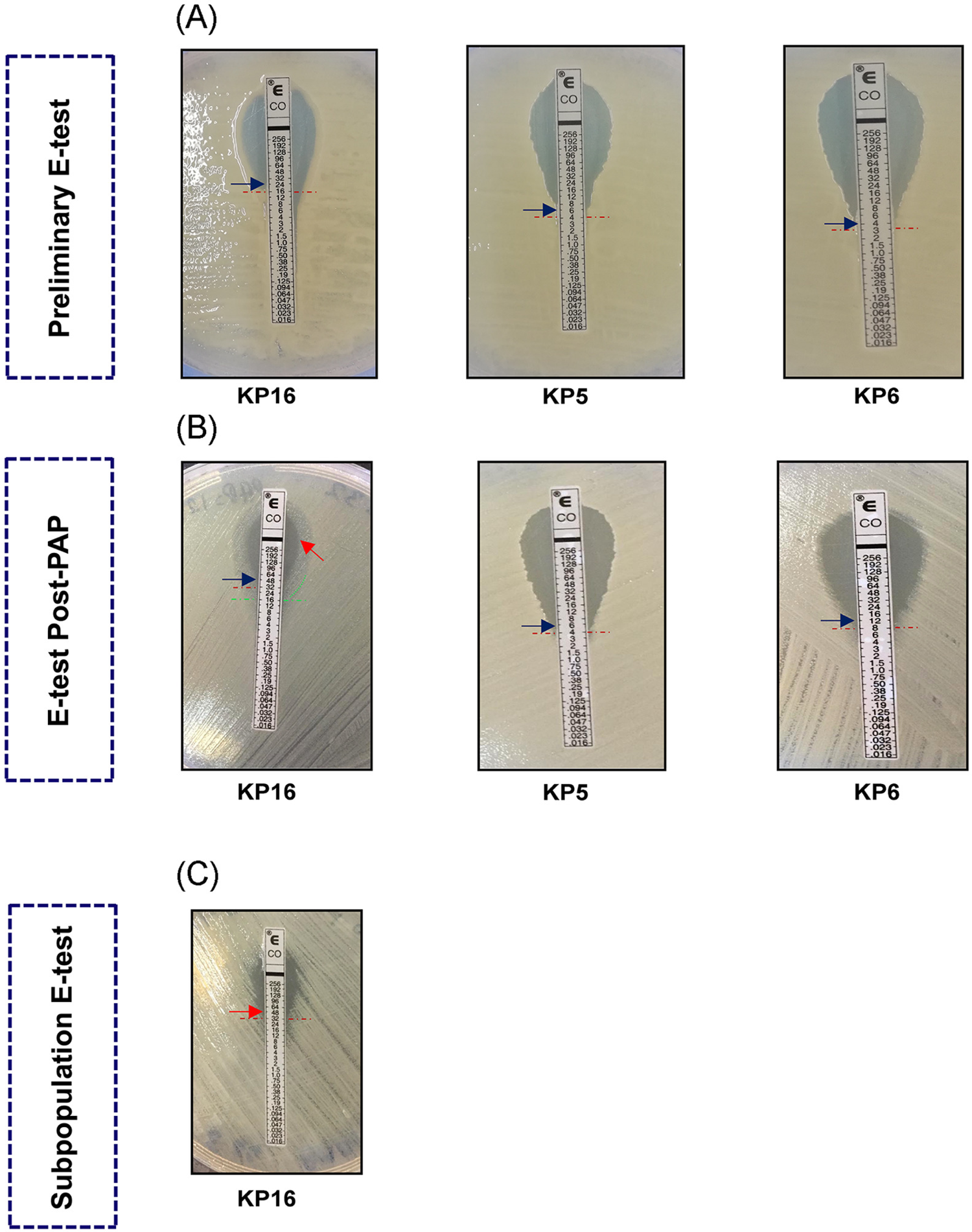
E-test antimicrobial susceptibility testing results of the highly colistin resistant isolates. A: initial testing of KP5, KP6, and KP16; B: E-test Post-PAP results; C: Subpopulation E-test.

**Figure 4:**
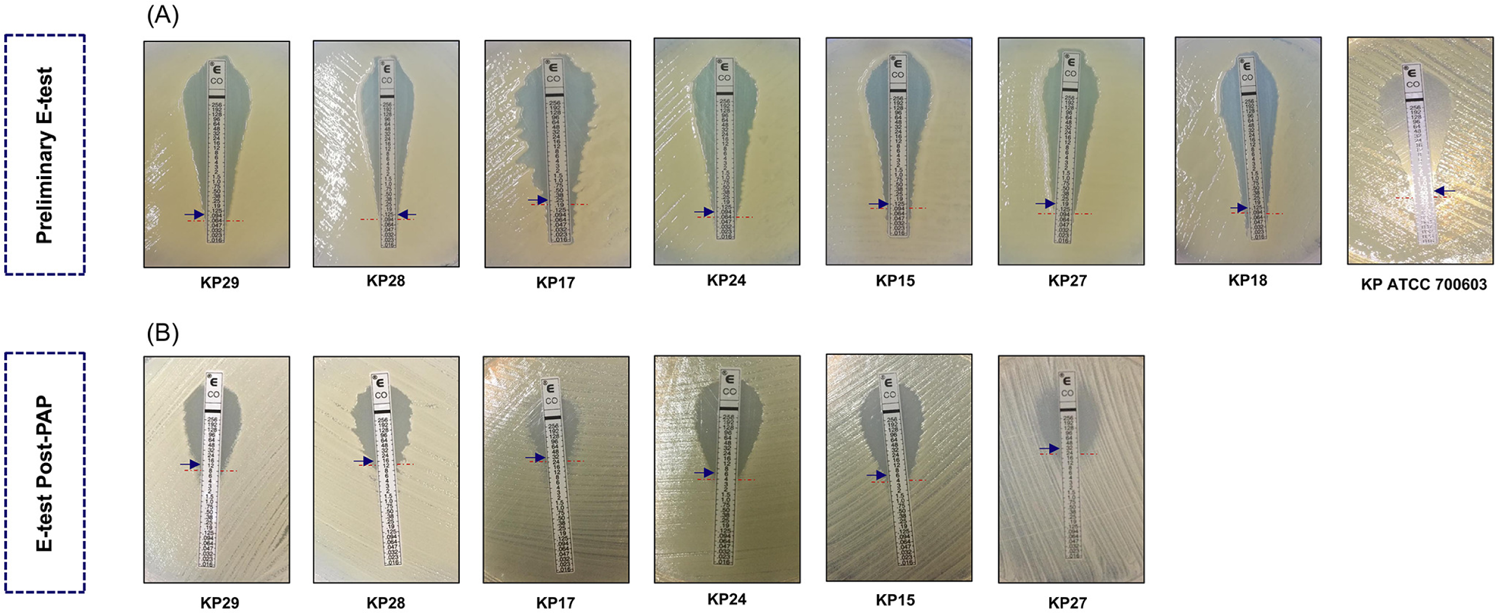
E-test antimicrobial susceptibility testing results of the highly colistin resistant isolates. A: initial testing of KP5, KP6, and KP16; B: E-test Post-PAP.

**Figure 5:**
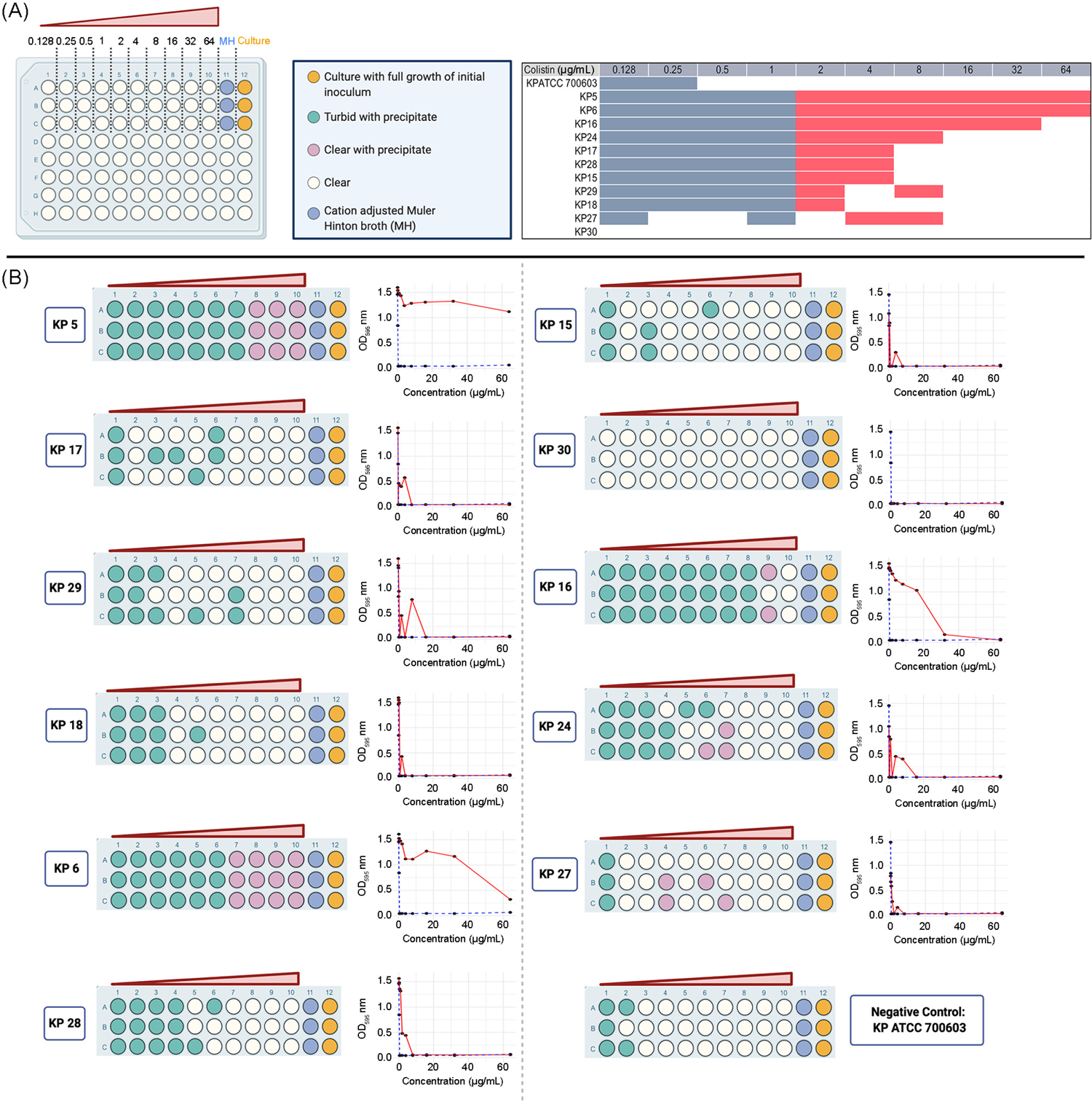
Broth Microdilution results. A: color coding representing the observed turbidity within the 96-well plate after BMD testing for 24 h. The table represents the MIC cutoff value with red bars ≥2 µg/mL; B: graphical representation of the phenomenon of heteroresistance and turbidity within each well. Optical density OD_595_ graphs are represented for each tested strain.

To further assess the stability of the heteroresistant phenotype, we aliquoted 1 μL from each well from the 24 h incubated BMD 96-well plate on colistin free-medium and incubated it at 37 °C for 16-24 h. The obtained *K. pneumoniae* colonies from the highest determined BMD MIC values were subjected to population analysis profiling (PAP). The PAP results showed an overall increase in the survival of the resistant subpopulations (Figure 6). KP16, which had the highest BMD MIC (64 µg/mL), showed normal growth pattern on PAP agar plates with 128 µg/mL of colistin. Similarly, KP17 and KP28 with BMD MIC of 4 µg/mL grew on PAP agar plates even after 4-fold increase in colistin concertation. KP24 and KP29 both had BMD MIC of 16 µg/mL but showed a 2-fold increase with the PAP assay. A similar 2-fold MIC increase was seen in KP15 and KP18. Only KP27 had the same MIC when tested first through BMD and after being exposed to colistin through PAP assay.

**Figure 6:**
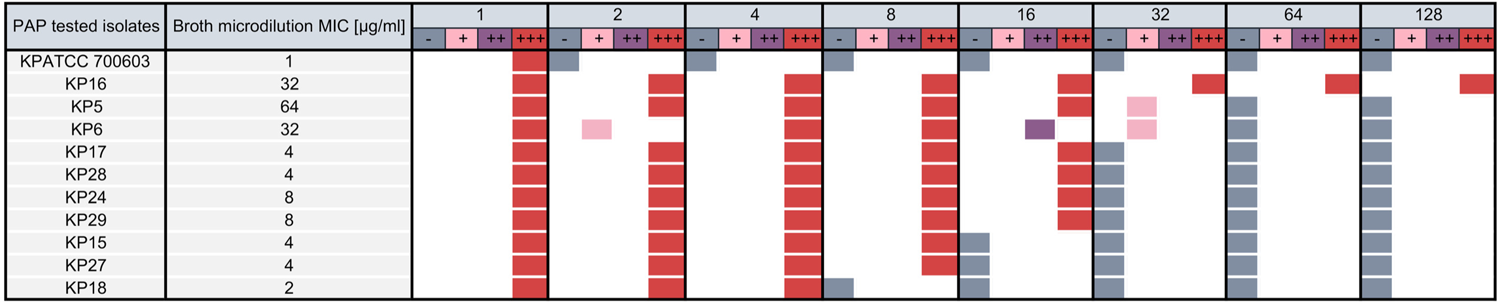
Population analysis profiling (PAP) of *K. pneumoniae* post exposure to colistin. BMD determined MIC (µg/mL) are listed. Colistin concentration with two-fold dilutions (128 µg/mL to 1 µg/mL). −: indicates no growth; +: indicates very few colonies; ++: indicates few colonies; +++: indicates complete growth of colonies.

To determine if the observed post-PAP higher MIC values is not due to a reversible phenotype, we subjected the recovered colonies from the highest PAP dilution MIC to a second E-test assay. A complex colistin resistance behavior was observed within the resistance subpopulation through E-test post-PAP results (Figure 3-B, and Figure 4-B). Interestingly, KP16 E-test post-PAP showed double halo distribution across the inhibition zone. Double MIC was 24 µg/mL for the heterogenous colistin resistant population and 46 µg/mL for the resistant sub-population (Figure 3-B). A third exposure of the recovered sub-population to colistin using the E-test (Figure 3-C) did not change the MIC value. Only KP5 had the same MIC 6 µg/mL after preliminary and post-PAP E-test, while KP16 post-PAP E-test MIC value was 3 folds higher (Figure 3-B). KP15, KP17, KP18, KP24, KP27, KP28, and KP29 showed several fold increases of the MIC following PAP (Figure 4-B).

### *K. pneumoniae* lipid A profiling through MALDI-TOF MS

All *K. pneumoniae* isolates were subjected to optimized acetic acid hydrolysis to extract and characterize the membrane lipids. The best output was obtained using 1% acetic acid solvent, while boiling at 100 °C for 1 h. Isolates KP5, KP6, and KP16 were colistin resistant according to the preliminary E-test. These isolates showed ions at m/z 1840 and 1824 representing bisphosphorylated and hexaacylated lipid A with or without hydroxylation, at position C’-2 of the fatty acyl chain, respectively. Ions at m/z 2063 were the outcome of palmitolation (-C-16) of the C-1 acyl-oxo-acyl chain at m/z 1840. On the other hand, m/z 2079 represented the palmitolation (-C-16) of the C-1 acyl-oxoacyl chain m/z 1824.

Moreover, KP5 with a colistin MIC value of > 64 µg/mL, had a modified lipid A structure detected through ion m/z 1892. A mass shift was generated because of lipid A modification which corresponded to L-Ara4N addition to the hexa-acylated lipid A structure m/z 1840 at C’-1 phosphate group, while losing a phosphate group at C’-1. The shift in mass could also be justified by the m/z 131 (equivalent to L-Ara4N) shift from m/z 1761 to m/z 1892. Additionally, ion at m/z 1956 represented a minor peak with a mass shift m/z 131 from m/z 1824 equivalent to glycosylation (L-Ara4N) of C’-1 phosphate group (Figure 7).

**Figure 7:**
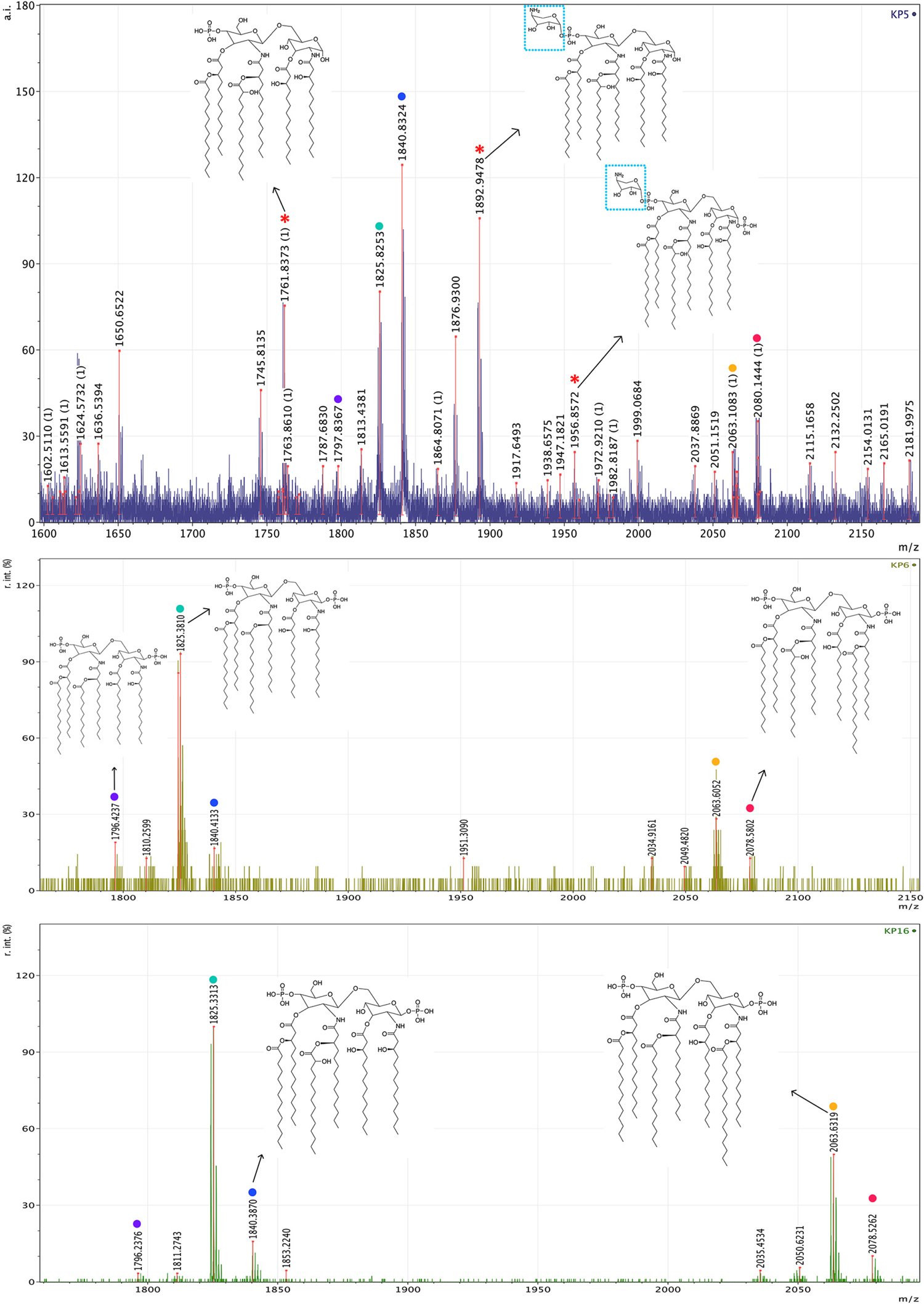
Lipid A ions detected in the negative reflectron mode from three colistin resistant *K. pneumoniae*. Each colored dot represents a sheared lipid A m/z value across KP5, KP6, and KP16; *: represents a modified lipid A structure distinct for KP5 isolate.

The remaining (8/11) isolates were profiled through MALDI-TOF MS and had their spectrum intensity normalized against lipid A standard. The results were like the ones obtained with respect to the m/z: 1840,1824, 2063 and 2079 (Figure 8-9).

**Figure 8:**
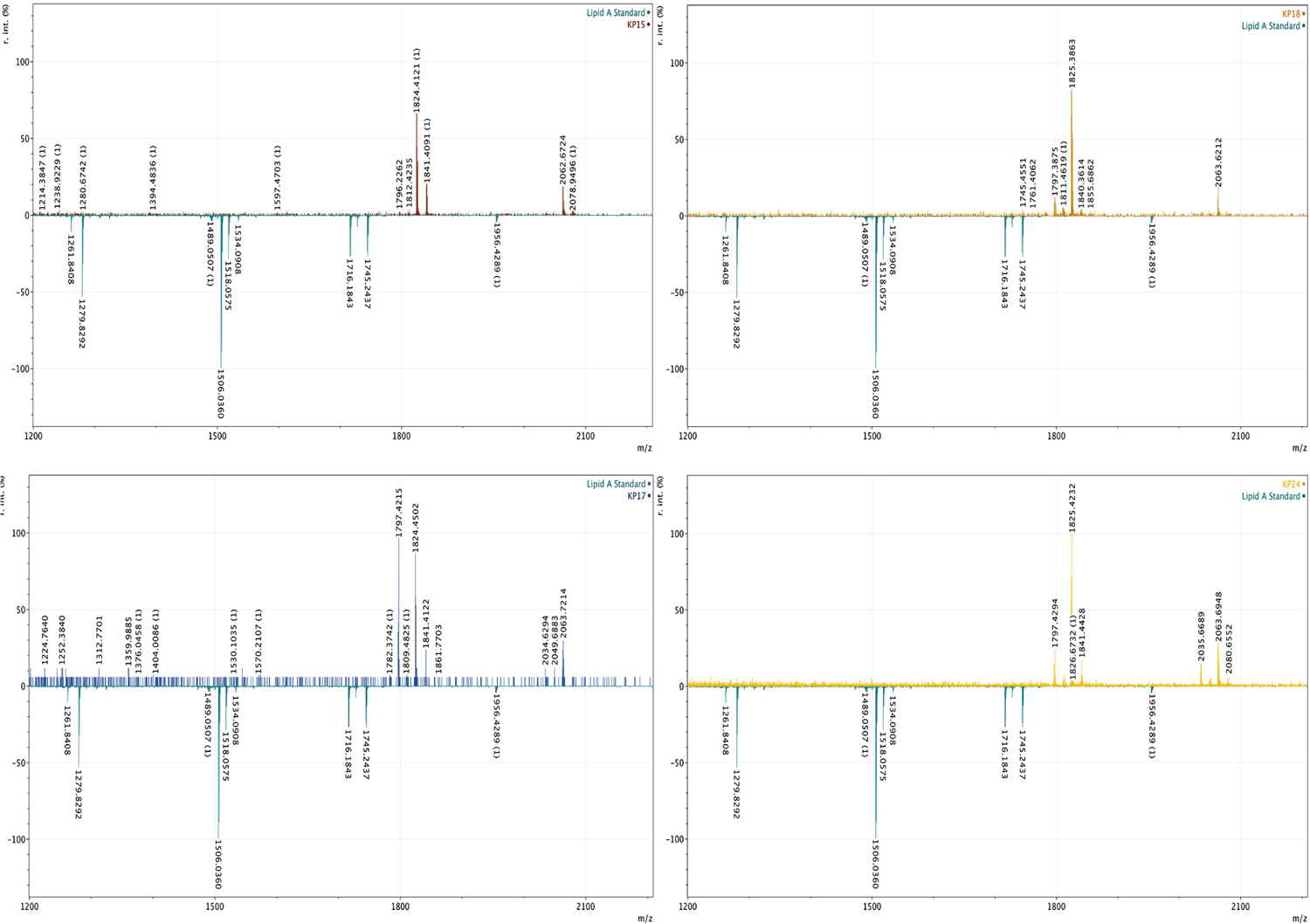
Lipid A profiling of isolates KP 15, KP18, KP17, KP24.

**Figure 9:**
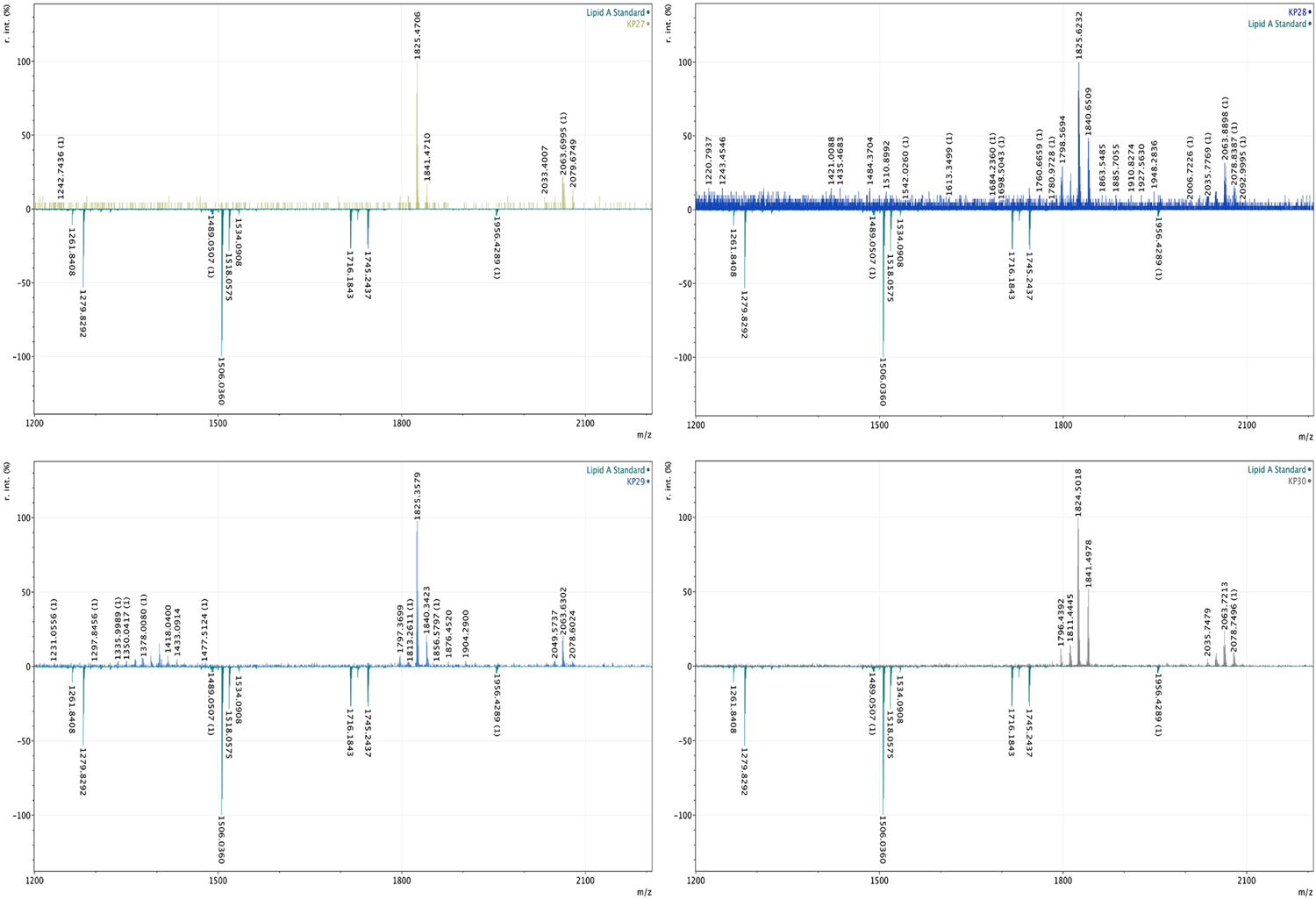
Lipid A profiling of isolates KP 27, KP28, KP29, KP30.

### *mgrB* locus identification and genetic environment

Upon analyzing the genetic environment surrounding the *mgr*B gene locus in the entire 32 set of isolates previously published by Arabaghian et al. 2019 (33), only three isolates (KP5, KP6, and KP16) displayed structural alterations in *mgr*B, resulting in the production of a dysfunctional gene product. This was further supported by the identification of two distinct types of insertion sequences, IS1R in KP5 and KP6 and a IS903B in KP16 belonging to the IS5 family. These insertion sequences divided the *mgr*B (Figure 10-11), into two fragments: *mgr*B-f1 (119 for KP5 and KP6 and 131 bp in KP16) and *mgr*B-f2 (85 bp for KP5 and KP6 and 64 bp for KP16). Additionally, the IS in KP5 and KP6 was bounded by direct repeats (GATTGCAC) (Figure 10), while a 126 bp repetitive element was detected in the IS903B-disrupted *mgrB* gene in KP16 (Figure 11). Sequence alignments of KP5 and KP6 and KP16 obtained by PCR amplification of the *mgrB* gene and whole-genome sequence data can be found in (Supplementary Figures 1 and 2), respectively.

**Figure 10:**
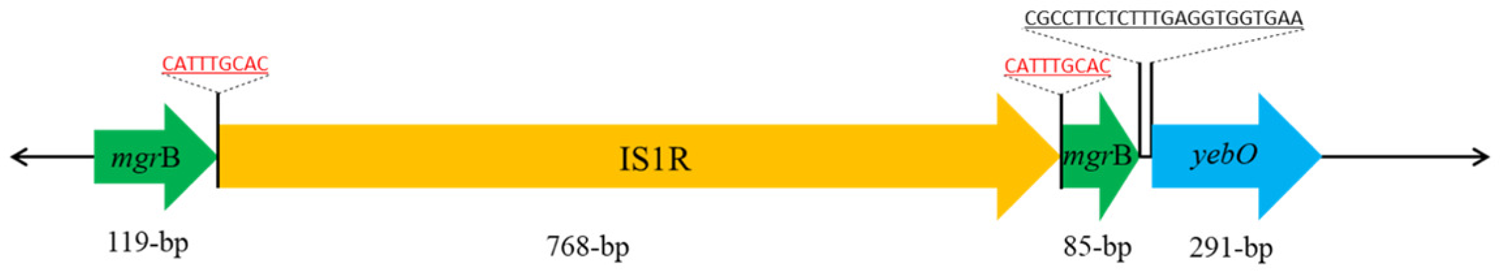
Schematic illustration of insertion sequence (IS1R) disrupting the *mgrB* in KP5 and KP6. The underlined red nucleotide sequence represents the direct repeats detected in KP5 and KP6.

**Figure 11:**
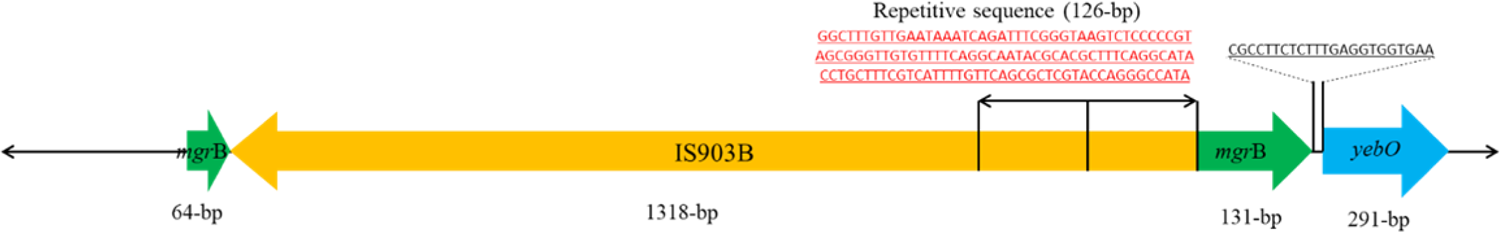
Schematic illustrations of insertion sequence (IS903B) disrupting the *mgrB* in KP16. The nucleotide sequence highlighted in red corresponds to a repetitive sequence (126 bp X2) within the insertion sequence.

### TCS amino acid substitutions and deletions

Various mutations were detected in TCSs prior to *in vitro* colistin exposure. An amino acid deletion in PhoQ (ΔM1 – L4) was observed in both KP5 and KP16, and other substitutions and deletions that could be directly linked to colistin resistance (Table 1).

**Table 1:**
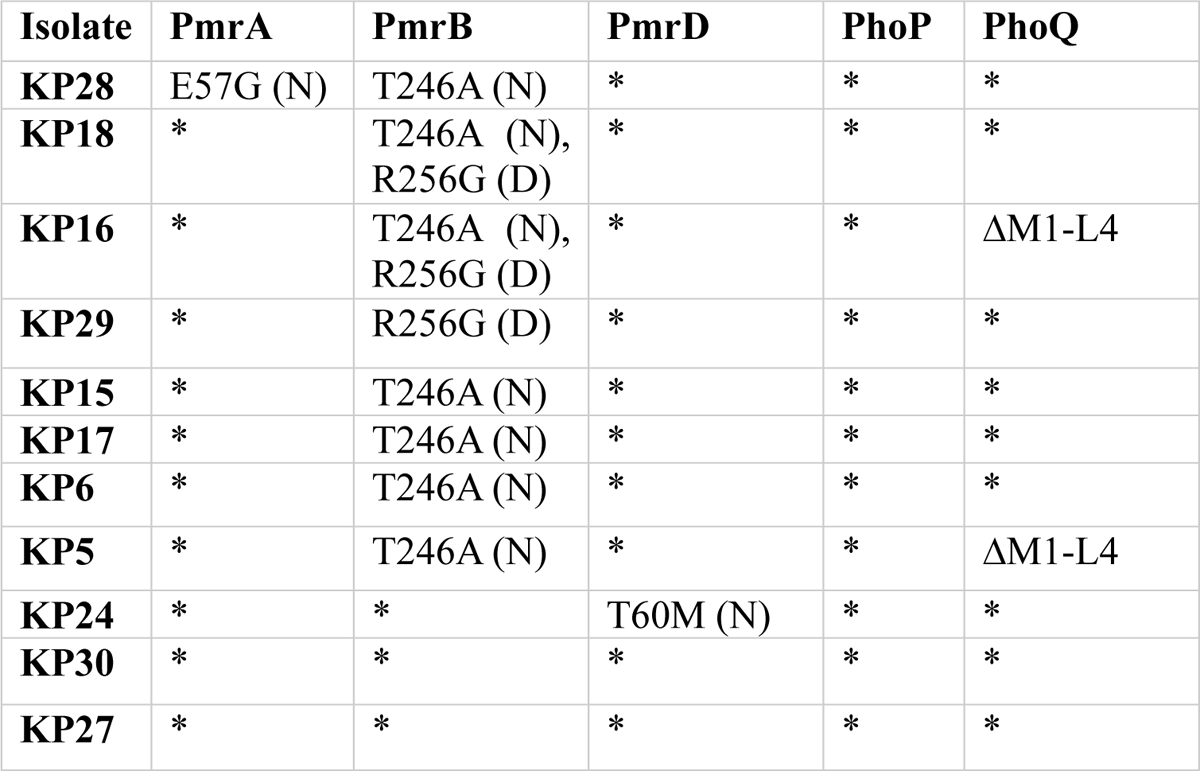
Features of *mgrB* and amino acid substitutions in the TCSs. *: identical to MGH 78578 sequence; Δ: deletion; (N): neutral effect of substitution on the protein; (D): deleterious effect of substitution on the protein.

| Isolate | PmrA | PmrB | PmrD | PhoP | PhoQ |
| --- | --- | --- | --- | --- | --- |
| **KP28** | E57G (N) | T246A (N) | * | * | * |
| **KP18** | * | T246A (N),<br>R256G (D) | * | * | * |
| **KP16** | * | T246A (N),<br>R256G (D) | * | * | ΔM1-L4 |
| **KP29** | * | R256G (D) | * | * | * |
| **KP15** | * | T246A (N) | * | * | * |
| **KP17** | * | T246A (N) | * | * | * |
| **KP6** | * | T246A (N) | * | * | * |
| **KP5** | * | T246A (N) | * | * | ΔM1-L4 |
| **KP24** | * | * | T60M (N) | * | * |
| **KP30** | * | * | * | * | * |
| **KP27** | * | * | * | * | * |

### Other known implicated colistin resistance genes

After conducting an extensive literature review, 29 genes including the TCSs mentioned earlier were found to be key components in the modification of LPS (Supplementary Table 1). Accordingly, those genes were examined within the 11 genomes understudy and were compared against the previously characterized *K. pneumoniae* isolates by Arabaghian et al 2019 (33). Utilizing a combination of genome annotation and blast tools, key observations were made regarding the annotations and nomenclature of these genes, with some genes having incomplete or alternative names (synonyms) (Supplementary Table 1). The *mcr* gene was not detected in any of the isolates understudy, while the *crr*B was absent in some of the 11 tested isolates (Supplementary Table 1). Importantly, our analysis of the amino acid sequence profiles did not reveal any specific mutations that marked resistance/heteroresistance. Further detailed amino acid substitutions can be found in (Supplementary Table 1).

## Discussion

We have previously studied the genetic content and molecular characteristics of 32 carbapenem resistant *K. pneumoniae* and two *K. quasipneumoniae* clinical isolates from Lebanon (33). In this work, we aimed at studying the susceptibility of the previously characterized *K. pneumoniae* to colistin followed by determining the underlying colistin resistance mechanisms leading to the complex phenotypes obtained post-colistin exposure. The main findings of this follow-up study reflected on the phenotypic diversity of colistin heterogeneous populations. Transient exposure to colistin through BMD and E-test assays facilitated the isolation of colistin resistant subpopulations with very high MIC values. Our results also revealed differences in resistance profiles obtained using colistin E-test versus BMD methods. We have shown that (7/11) of the isolates that were thought to be colistin sensitive through preliminary E-test assay had established high level of resistance to colistin. The resistance was likely due to chromosomal SNPs. On the other hand, the colistin resistant isolates (KP5, KP6, and KP16), were all negative for *mcr*, exhibited a heterogeneous response to colistin, or had a transient adaptive resistance following three phases of exposure assays.

Heteroresistant subpopulations could be detected using the E-test generated zone of inhibition (ellipse-shape), but this should be cautiously considered. Band et al. (15) previously showed that colistin heteroresistant is largely undetected in the clinic. Alternative approaches such as disk diffusion and gradient diffusion were also unreliable, mainly due to the cationic nature and large size of polymyxins (47). Heteroresistance was linked however, to uninterpreted MIC values associated with more than one skip-well/tested concentration, which could be demonstrated when detecting wells with no growth while being observed in ones with higher concentrations of the antibiotic (48). PAP provided a better insight and helped in revealing changes in susceptibility upon exposure.

Silva et al. (49) showed using PAP that *K. pneumoniae* manifested heteroresistance to colistin only when grown in biofilm arrangements, and a resistant sub-population was identified. Resistance to colistin in *K. pneumoniae* could also result from the disruption of the *mgrB* gene by ISs or through missense or nonsense mutations leading to a stop codon (13, 31, 48). We detected insertional inactivation of *mgrB* gene in the three colistin resistant isolates (KP5, KP6, and KP16). KP5 and KP6 both belonged to ST1799, shared the same capsular type K16, and had IS1R mediated *mgrB* disruption. IS1R belongs to IS1 family and was reported in colistin resistant *K. pneumoniae* isolated from Greece (50), Moreover, KP16 which belonged to sequence type ST147 and capsular type K64, had another IS element IS903B, a member of IS5 family. The similarity between the IS5 element inserted into *mgrB* and the IS5 element present on *K. pneumoniae bla*_KPC_ encoding plasmids revealed possible mobilization of the IS5 elements from plasmids into the *mgrB* gene (31). Furthermore, Yang et al. (51) showed that for the plasmid carrying IS903B, colistin induced stress was responsible for IS mobilization.

Various point mutations leading to amino acid substitutions in PmrA, PmrB, PhoP, PhoQ, and PmrD were also detected in this study. Multiple amino acid substitutions observed were also previously reported (52, 53) and linked to colistin resistance in *K. pneumoniae* and could be involved in inducing resistance, but that requires further verification and which was beyond the scope of this study.

Only one R256G substitution in PmrB coding region in KP18 and KP16 was predicted to have deleterious effect on the protein function. R256G substitution was also reported in a study from Iran by Haeili et al. (52) in colistin resistant *K. pneumoniae* carrying a wild type *mgrB* gene. The authors however, showed that complementation with the wild type PmrB did not restore colistin susceptibility; colistin MIC before and after complementation remained equal to 128 mg/L (52). Additionally, T246A substitution in PmrB was detected in colistin resistant *K. pneumoniae* recovered from a Tunisian Teaching Hospital (53) which we also found in KP28, KP18, KP16, KP15, KP17, KP6, and KP5. On the other hand, PmrA E57G substitution in KP28 and PmrD T60M substitution in KP24 had a neutral effect and were not previously reported. Interestingly, a PhoQ deletion ΔM1-L4 was observed for the first time in the highly resistant isolates, KP5 and KP16. This deletion to our knowledge was not reported previously. However, a different PhoQ deletion ΔK2-L6 was detected in five colistin resistant *K. pneumoniae* belonging to ST101 in Tunisia (53) (Table 1).

We also screened for synonymous and non-synonymous mutations in genes linked to outer membrane biosynthesis. *arnT* was linked to modifications mediating colistin resistance through L-Ara4N in *Salmonella* Typhimurium. We detected a non-synonymous mutation leading to L296V substitution in *arnT*. Nonetheless, cation transporter genes (*mntH* and *nhaR*), MFS family efflux pumps (*yceE*), efflux pump (*acrA, acrB*), and LPS biosynthesis gene (*lpxD*) were found to only harbor synonymous mutations. Very few studies stressed on the role of efflux pumps in polymyxin resistance. *acrA* and *acrB* code for the AcrAB-TolC efflux pump in *K. pneumoniae* which was associated with resistance to cationic antimicrobial peptides and fluoroquinolones. Isolates with mutations in *acrB* were according to (54) significantly more susceptible to polymyxin B than the wild-type strains. We only detected synonymous mutations in AcrA encoding genes in all tested isolates (Supplementary Table 1), and thus we excluded this system as a potential candidate mediating resistance. Future studies of manual curation and experimental validation may be needed to validate the computationally inferred effects of the detected amino acid substitutions and deletions.

Colistin resistance arises from chemical modifications to the lipid A portion of the LPS. We used MALDI-TOF-based approaches to detect and verify colistin resistance. LPS extraction and detection procedures are variable and dependent on the studied pathogen. An optimized workflow to obtain lipid A from the outer membrane of *E. coli* and *K. pneumoniae* was established in this study. A schematic representation of the extraction procedure is presented in (Figure 1). KP5, colistin resistant isolate, had the L-Ara4N peak at m/z 1892, which to the best of our knowledge was not previously described. It corresponded to L-Ara4N addition to the hexa-acylated lipid A structure m/z 1840 at C’-1 phosphate group and to loss of C-1 phosphate group. The latter could be linked to the acidic conditions used in the extraction procedure (55). In addition, a lower intensity peak at m/z 1955, linked to L-Ara4N modified lipid A was detected in KP5 and was previously reported in colistin resistant *K. pneumoniae* (56). In contrast, KP6 and KP16 which should also exhibit lipid A modification being highly resistant to colistin, had a native lipid A peak distribution. A finding that merits further studies and which was not within the scope of this work. The results obtained from the remaining colistin heteroresistant isolates and one colistin sensitive *K. pneumoniae* were congruent with the previous published work related to colistin sensitive *K. pneumoniae* (56).

In conclusion, *K. pneumoniae* are opportunistic pathogens which can cause different types of healthcare-associated infections and are difficult to treat. Colistin resistance is known to be inducible during colistin treatment and can be also caused through mutations and genetic alterations in regulatory networks controlling chemical modifications of the lipid A. Interruptions in *mgrB* that increase expression of the *pmrHFIJKLM* operon are major mechanisms contributing to colistin resistance. The dissemination of IS elements that transpose into the *mgrB* gene was an important mechanism mediating the observed colistin resistance in this study and was linked to the observed elevated MICs. Furthermore, resistant subpopulations with chromosomal alterations can become dominant during a colistin selection pressure and facilitate the emergence and spread of resistance leading to treatment failure and is a serious clinical concern that merits further studies.

---

**Supplementary Figure 1:** Multiple sequence alignment for *mgrB* genetic region. KP5 and KP6 sequences were retrieved from whole genome sequencing contigs, and Sanger Sequencing. IS1R insertion sequence and IS1D insertion sequence were also aligned to reflect the areas on insertion.

## References

1. Rochford C, Sridhar D, Woods N, Saleh Z, Hartenstein L, Ahlawat H, Whiting E, Dybul M, Cars O, Goosby E, Cassels A, Velasquez G, Hoffman S, Baris E, Wadsworth J, Gyansa-Lutterodt M, Davies S. 2018. Global governance of antimicrobial resistance. Lancet 391:1976–1978.

2. O’Neill J. 2014. Antimicrobial resistance: tackling a crisis for the health and wealth of nations. https://www.who.int/news/item/29-04-2019-new-report-calls-for-urgent-action-to-avert-antimicrobial-resistance-crisis.

3. Falagas ME, Kasiakou SK. 2005. Colistin: the revival of polymyxins for the management of multidrug-resistant gram-negative bacterial infections. Clin Infect Dis 40:1333–41.

4. Paczosa MK, Mecsas J. 2016. *Klebsiella pneumoniae*: going on the offense with a strong defense. Microbiol Mol Biol Rev 80:629–61.

5. Rojas LJ, Salim M, Cober E, Richter SS, Perez F, Salata RA, Kalayjian RC, Watkins RR, Marshall S, Rudin SD, Domitrovic TN, Hujer AM, Hujer KM, Doi Y, Kaye KS, Evans S, Fowler VG, Jr., Bonomo RA, van Duin D. 2017. Colistin resistance in carbapenem-resistant *Klebsiella pneumoniae*: laboratory detection and impact on mortality. Clin Infect Dis 64:711–718.

6. Poirel L, Jayol A, Nordmann P. 2017. Polymyxins: antibacterial activity, susceptibility testing, and resistance mechanisms encoded by plasmids or chromosomes. Clin Microbiol Rev 30:557–596.

7. Liu YY, Wang Y, Walsh TR, Yi LX, Zhang R, Spencer J, Doi Y, Tian G, Dong B, Huang X, Yu LF, Gu D, Ren H, Chen X, Lv L, He D, Zhou H, Liang Z, Liu JH, Shen J. 2016. Emergence of plasmid-mediated colistin resistance mechanism MCR-1 in animals and human beings in China: a microbiological and molecular biological study. Lancet Infect Dis 16:161–8.

8. Gunn JS, Miller SI. 1996. PhoP-PhoQ activates transcription of *pmrAB*, encoding a two-component regulatory system involved in *Salmonella typhimurium* antimicrobial peptide resistance. J Bacteriol 178:6857–64.

9. Helander IM, Kato Y, Kilpeläinen I, Kostiainen R, Lindner B, Nummila K, Sugiyama T, Yokochi T. 1996. Characterization of lipopolysaccharides of polymyxin-resistant and polymyxin-sensitive *Klebsiella pneumoniae* O3. Eur J Biochem 237:272–8.

10. Mitrophanov AY, Jewett MW, Hadley TJ, Groisman EA. 2008. Evolution and Dynamics of Regulatory Architectures Controlling Polymyxin B Resistance in Enteric Bacteria. PLoS Genet 4:e1000233.

11. Nikaido H. 2003. Molecular basis of bacterial outer membrane permeability revisited. Microbiol Mol Biol Rev 67:593–656.

12. Lippa AM, Goulian M. 2009. Feedback inhibition in the PhoQ/PhoP signaling system by a membrane peptide. PLoS Genet 5:e1000788.

13. Cannatelli A, Giani T, D’Andrea MM, Di Pilato V, Arena F, Conte V, Tryfinopoulou K, Vatopoulos A, Rossolini GM. 2014. MgrB inactivation is a common mechanism of colistin resistance in KPC-producing *Klebsiella pneumoniae* of clinical origin. Antimicrob Agents Chemother 58:5696–703.

14. Cheng YH, Lin TL, Lin YT, Wang JT. 2016. Amino Acid Substitutions of CrrB Responsible for Resistance to Colistin through CrrC in *Klebsiella pneumoniae*. Antimicrob Agents Chemother 60:3709–16.

15. Band VI, Satola SW, Burd EM, Farley MM, Jacob JT, Weiss DS. 2018. Carbapenem-resistant *Klebsiella pneumoniae* exhibiting clinically undetected colistin heteroresistance leads to treatment failure in a murine model of infection. mBio 9:e02448–17.

16. Liao W, Lin J, Jia H, Zhou C, Zhang Y, Lin Y, Ye J, Cao J, Zhou T. 2020. Resistance and heteroresistance to colistin in *Escherichia coli* isolates from Wenzhou, China. Infect Drug Resist 13:3551–3561.

17. Falagas ME, Makris GC, Dimopoulos G, Matthaiou DK. 2008. Heteroresistance: a concern of increasing clinical significance? Clin Microbiol Infect 14:101–4.

18. El-Halfawy OM, Valvano MA. 2015. Antimicrobial heteroresistance: an emerging field in need of clarity. Clin Microbiol Rev 28:191–207.

19. Wright MS, Suzuki Y, Jones MB, Marshall SH, Rudin SD, van Duin D, Kaye K, Jacobs MR, Bonomo RA, Adams MD. 2015. Genomic and transcriptomic analyses of colistin-resistant clinical isolates of *Klebsiella pneumoniae* reveal multiple pathways of resistance. Antimicrob Agents Chemother 59:536–43.

20. Jayol A, Nordmann P, Brink A, Poirel L. 2015. Heteroresistance to colistin in *Klebsiella pneumoniae* associated with alterations in the PhoPQ regulatory system. Antimicrob Agents Chemother 59:2780–4.

21. CLSI. 2014. Performance standards for antimicrobial susceptibility testing; 24th informational supplement on Clinical and Laboratory Standards Institute, Wayne, PA. https://www.researchgate.net/file.PostFileLoader.html?id=59202a0696b7e4d462166956&assetKey=AS%3A496054988533760%401495280134033.

22. EUCAST. 2014. Breakpoint tables for interpretation of MICs and zone diameters, version 2.0, on European Committee on Antimicrobial Susceptibility Testing (EUCAST). http://www.eucast.org/fileadmin/src/media/PDFs/EUCAST_files/Breakpoint_tables/Breakpoint_table_v_2.0_120221.pdf.

23. Albiger B, Glasner C, Struelens MJ, Grundmann H, Monnet DL. 2015. Carbapenemase-producing Enterobacteriaceae in Europe: assessment by national experts from 38 countries, May 2015. Euro Surveill 20.

24. Arduino SM, Quiroga MP, Ramírez MS, Merkier AK, Errecalde L, Di Martino A, Smayevsky J, Kaufman S, Centrón D. 2012. Transposons and integrons in colistin-resistant clones of *Klebsiella pneumoniae* and *Acinetobacter baumannii* with epidemic or sporadic behaviour. J Med Microbiol 61:1417–1420.

25. Kontopoulou K, Protonotariou E, Vasilakos K, Kriti M, Koteli A, Antoniadou E, Sofianou D. 2010. Hospital outbreak caused by *Klebsiella pneumoniae* producing KPC-2 beta-lactamase resistant to colistin. J Hosp Infect 76:70–3.

26. Marchaim D, Chopra T, Pogue JM, Perez F, Hujer AM, Rudin S, Endimiani A, Navon-Venezia S, Hothi J, Slim J, Blunden C, Shango M, Lephart PR, Salimnia H, Reid D, Moshos J, Hafeez W, Bheemreddy S, Chen TY, Dhar S, Bonomo RA, Kaye KS. 2011. Outbreak of colistin-resistant, carbapenem-resistant *Klebsiella pneumoniae* in metropolitan Detroit, Michigan. Antimicrob Agents Chemother 55:593–9.

27. Bogdanovich T, Adams-Haduch JM, Tian GB, Nguyen MH, Kwak EJ, Muto CA, Doi Y. 2011. Colistin-resistant, Klebsiella pneumoniae carbapenemase (KPC)-producing Klebsiella pneumoniae belonging to the international epidemic clone ST258. Clin Infect Dis 53:373–6.

28. Mammina C, Bonura C, Di Bernardo F, Aleo A, Fasciana T, Sodano C, Saporito MA, Verde MS, Tetamo R, Palma DM. 2012. Ongoing spread of colistin-resistant *Klebsiella pneumoniae* in different wards of an acute general hospital, Italy, June to December 2011. Euro Surveill 17.

29. Berglund B, Hoang NTB, Tärnberg M, Le NK, Svartström O, Khu DTK, Nilsson M, Le HT, Welander J, Olson L, Larsson M, Nilsson LE, Hanberger H. 2018. Insertion sequence transpositions and point mutations in *mgrB* causing colistin resistance in a clinical strain of carbapenem-resistant *Klebsiella pneumoniae* from Vietnam. Int J Antimicrob Agents 51:789–793.

30. Salloum T, Panossian B, Bitar I, Hrabak J, Araj GF, Tokajian S. 2020. First report of plasmid-mediated colistin resistance *mcr-8.1* gene from a clinical *Klebsiella pneumoniae* isolate from Lebanon. Antimicrob Resist Infect Control 9, 94. 10.1186/s13756-020-00759-w31.

31. Cannatelli A, D’Andrea MM, Giani T, Di Pilato V, Arena F, Ambretti S, Gaibani P, Rossolini GM. 2013. In vivo emergence of colistin resistance in *Klebsiella pneumoniae* producing KPC-type carbapenemases mediated by insertional inactivation of the PhoQ/PhoP *mgrB* regulator. Antimicrob Agents Chemother 57:5521–6.

32. Choi Y, Chan AP. 2015. PROVEAN web server: a tool to predict the functional effect of amino acid substitutions and indels. Bioinform 31:2745–7.

33. Arabaghian H, Salloum T, Alousi S, Panossian B, Araj GF, Tokajian S. 2019. Molecular Characterization of Carbapenem Resistant *Klebsiella pneumoniae* and *Klebsiella quasipneumoniae* Isolated from Lebanon. Sci Rep 9:531.

34. Andrews S. 2010. FastQC: A Quality Control Tool for High Throughput Sequence Dat. http://www.bioinformatics.babraham.ac.uk/projects/fastqc/.

35. Bolger AM, Lohse M, Usadel B. 2014. Trimmomatic: a flexible trimmer for Illumina sequence data. Bioinform 30:2114–20.

36. Bankevich A, Nurk S, Antipov D, Gurevich AA, Dvorkin M, Kulikov AS, Lesin VM, Nikolenko SI, Pham S, Prjibelski AD, Pyshkin AV, Sirotkin AV, Vyahhi N, Tesler G, Alekseyev MA, Pevzner PA. 2012. SPAdes: a new genome assembly algorithm and its applications to single-cell sequencing. J Comput Biol 19:455–77.

37. Gurevich A, Saveliev V, Vyahhi N, Tesler G. 2013. QUAST: quality assessment tool for genome assemblies. Bioinform 29:1072–5.

38. Wood DE, Lu J, Langmead B. 2019. Improved metagenomic analysis with Kraken 2. Genome Biol 20.

39. Wick RR, Schultz MB, Zobel J, Holt KE. 2015. Bandage: interactive visualization of de novo genome assemblies. Bioinform 31:3350–2.

40. Lakin SM, Kuhnle A, Alipanahi B, Noyes NR, Dean C, Muggli M, Raymond R, Abdo Z, Prosperi M, Belk KE, Morley PS, Boucher C. 2019. Hierarchical Hidden Markov models enable accurate and diverse detection of antimicrobial resistance sequences. Commun Biol 2:294.

41. Seemann T. 2014. Prokka: rapid prokaryotic genome annotation. Bioinform 30:2068–9.

42. Alcock BP, Raphenya AR, Lau TTY, Tsang KK, Bouchard M, Edalatmand A, Huynh W, Nguyen A-LV, Cheng AA, Liu S, Min SY, Miroshnichenko A, Tran H-K, Werfalli RE, Nasir JA, Oloni M, Speicher DJ, Florescu A, Singh B, Faltyn M, Hernandez-Koutoucheva A, Sharma AN, Bordeleau E, Pawlowski AC, Zubyk HL, Dooley D, Griffiths E, Maguire F, Winsor GL, Beiko RG, Brinkman FSL, Hsiao WWL, Domselaar GV, McArthur AG. 2019. CARD 2020: antibiotic resistome surveillance with the comprehensive antibiotic resistance database. Nucleic Acids Res 48:D517–D525.

43. Bonin N, Doster E, Worley H, Pinnell LJ, Bravo JE, Ferm P, Marini S, Prosperi M, Noyes N, Morley PS, Boucher C. 2022. MEGARes and AMR++, v3.0: an updated comprehensive database of antimicrobial resistance determinants and an improved software pipeline for classification using high-throughput sequencing. Nucleic Acids Res 51:D744–D752.

44. Katoh K, Rozewicki J, Yamada KD. 2019. MAFFT online service: multiple sequence alignment, interactive sequence choice and visualization. Brief Bioinform 20:1160–1166.

45. Guindon S, Dufayard J-F, Lefort V, Anisimova M, Hordijk W, Gascuel O. 2010. New Algorithms and Methods to Estimate Maximum-Likelihood Phylogenies: Assessing the Performance of PhyML 3.0. Syst Biol 59:307–321.

46. Caroff M, Tacken A, Szabó L. 1988. Detergent-accelerated hydrolysis of bacterial endotoxins and determination of the anomeric configuration of the glycosyl phosphate present in the “isolated lipid A” fragment of the *Bordetella pertussis* endotoxin. Carbohydr Res 175:273–82.

47. EUCAST. 2016. Recommendations for MIC determination of colistin (polymyxin E) as recommended by the joint CLSI-EUCAST Polymyxin Breakpoints Working Group. http://www.eucast.org/guidance_documents/.

48. Olaitan AO, Diene SM, Kempf M, Berrazeg M, Bakour S, Gupta SK, Thongmalayvong B, Akkhavong K, Somphavong S, Paboriboune P, Chaisiri K, Komalamisra C, Adelowo OO, Fagade OE, Banjo OA, Oke AJ, Adler A, Assous MV, Morand S, Raoult D, Rolain JM. 2014. Worldwide emergence of colistin resistance in *Klebsiella pneumoniae* from healthy humans and patients in Lao PDR, Thailand, Israel, Nigeria and France owing to inactivation of the PhoP/PhoQ regulator *mgrB*: an epidemiological and molecular study. Int J Antimicrob Agents 44:500–7.

49. Silva A, Sousa AM, Alves D, Lourenço A, Pereira MO. 2016. Heteroresistance to colistin in *Klebsiella pneumoniae* is triggered by small colony variants sub-populations within biofilms. Pathog Dis 74.

50. Hamel M, Chatzipanagiotou S, Hadjadj L, Petinaki E, Papagianni S, Charalampaki N, Tsiplakou S, Papaioannou V, Skarmoutsou N, Spiliopoulou I. 2020. Inactivation of *mgrB* gene regulator and resistance to colistin is becoming endemic in carbapenem-resistant *Klebsiella pneumoniae* in Greece: A nationwide study from 2014 to 2017. Int J Antimicrob Agents 55:105930.

51. Yang TY, Wang SF, Lin JE, Griffith BTS, Lian SH, Hong ZD, Lin L, Lu PL, Tseng SP. 2020. Contributions of insertion sequences conferring colistin resistance in *Klebsiella pneumoniae*. Int J Antimicrob Agents 55:105894.

52. Haeili M, Javani A, Moradi J, Jafari Z, Feizabadi MM, Babaei E. 2017. MgrB alterations mediate colistin resistance in *Klebsiella pneumoniae* isolates from Iran. Front Microbiol 8:2470.

53. Jaidane N, Bonnin RA, Mansour W, Girlich D, Creton E, Cotellon G, Chaouch C, Boujaafar N, Bouallegue O, Naas T. 2018. Genomic insights into colistin-resistant *Klebsiella pneumoniae* from a Tunisian teaching hospital. Antimicrob Agents Chemother 62(2):e01601–17. doi: 10.1128/AAC.01601-17.54.

54. Padilla E, Llobet E, Doménech-Sánchez A, Martínez-Martínez L, Bengoechea JA, Albertí S. 2010. *Klebsiella pneumoniae* AcrAB efflux pump contributes to antimicrobial resistance and virulence. Antimicrob Agents Chemother 54:177–83.

55. Karibian D, Deprun C, Caroff M. 1995. Use of plasma desorption mass spectrometry in structural analysis of endotoxins: effects on lipid A of different acid treatments. Prog Clin Biol Res 392:103–11.

56. Leung LM, Cooper VS, Rasko DA, Guo Q, Pacey MP, McElheny CL, Mettus RT, Yoon SH, Goodlett DR, Ernst RK, Doi Y. 2017. Structural modification of LPS in colistin-resistant, KPC-producing *Klebsiella pneumoniae*. J Antimicrob Chemother 72:3035–3042.

